# A Corollary Discharge Circuit in Human Speech

**DOI:** 10.1101/2022.09.12.507590

**Authors:** Amirhossein Khalilian-Gourtani, Ran Wang, Xupeng Chen, Leyao Yu, Patricia Dugan, Daniel Friedman, Werner Doyle, Orrin Devinsky, Yao Wang, Adeen Flinker

## Abstract

When we vocalize, our brain distinguishes self-generated sounds from external ones. A corollary discharge signal supports this function in animals, however, in humans its exact origin and temporal dynamics remain unknown. We report Electrocorticographic (ECoG) recordings in neurosurgical patients and a novel connectivity approach based on Granger-causality that reveals major neural communications. We find a reproducible source for corollary discharge across multiple speech production paradigms localized to ventral speech motor cortex before speech articulation. The uncovered discharge predicts the degree of auditory cortex suppression during speech, its well-documented consequence. These results reveal the human corollary discharge source and timing with far-reaching implication for speech motor-control as well as auditory hallucinations in human psychosis.

**Significance statement:** How do organisms dissociate self-generated sounds from external ones? A fundamental brain circuit across animals addresses this question by transmitting a blueprint of the motor signal to sensory cortices, referred to as a corollary discharge. However, in humans and non-human primates auditory system, the evidence supporting this circuit has been limited to its direct consequence, auditory suppression. Furthermore, an impaired corollary discharge circuit in humans can lead to auditory hallucinations. While hypothesized to originate in the frontal cortex, direct evidence localizing the source and timing of an auditory corollary discharge is lacking in humans. Leveraging rare human neurosurgical recordings combined with connectivity techniques, we elucidate the exact source and dynamics of the corollary discharge signal in human speech.

**One-sentence summary:** We reveal the source and timing of a corollary discharge from speech motor cortex onto auditory cortex in human speech.

## 1 Introduction

How does the brain dissociate self-generated stimuli from external ones? Any motor act directly activates associated sensory systems. This constant flow of sensory information, while useful as feedback, can desensitize the sensory system or be confused with external sensations (*1, 2*). A fundamental brain circuit solves this problem by transmitting a blueprint of the motor signal to sensory cortices, referred to as corollary discharge (*3*). The corollary discharge signals are established in many species and multiple sensory modalities (*1, 2, 4–6*). In the human auditory system, a corollary discharge is hypothesized to increase sensitivity to self-generated speech during production (*7–9*) and when impaired can lead to auditory hallucinations (*10, 11*).

Corollary discharge signals decrease the sensory processing load and increase sensitivity during vocalization by suppressing sensory cortices (*12, 13*). For example, an inter-neuron in the cricket motor system inhibits their auditory system during loud chirping to avoid desensitization (*2*). In non-human primates, vocalization suppresses auditory neurons, supporting an auditory corollary discharge circuit (*12*). Similarly, self-produced human speech suppresses auditory cortex (*14–18*) and schizophrenia patients with auditory hallucinations have impaired suppression (*10, 11*). In primates and humans, the source of the corollary discharge signal and its dynamics remain virtually unknown (*14, 19*).

Recent neuroscience approaches focus on the study of connectivity and information flow between cortical regions (*20–22*). However, many leverage non-invasive neuroimaging data with a limited temporal (i.e. fMRI) or spatial (i.e M/EEG) resolution and typically do not assess the directionality of information flow (e.g. functional connectivity). Here, we leverage rare human neurosurgical recordings from motor and auditory related cortical sites during speech production tasks providing both a high spatial and temporal resolution. Leveraging directed connectivity approaches based on Granger-causality combined with unsupervised learning we identify the source, target, and temporal dynamics of information flow across cortices. A discharge signal before speech onset is transmitted from ventral motor cortex to auditory cortex, and is reproducible across multiple speech tasks and patients. This directed signal from motor cortex predicts the degree of suppression across auditory sites, providing the first direct evidence for a corollary discharge signal and its temporal dynamics in humans.

## 2 Results

To investigate the corollary discharge (CD) signal in human speech, we employed a paradigm that directly measures the CD signal’s outcome, i.e., speech induced auditory suppression (*14*). We acquired electrocorticographic (ECoG) recordings from eight neurosurgical patients while they performed an auditory word repetition task. We focused on high-gamma broadband (70-150 Hz), a marker of neural activity. We first established the temporal and spatial neural recruitment during auditory perception (Fig. 1A) and production (Fig. 1B). Neural activity commenced in auditory cortices (i.e. superior temporal gyrus; STG) shortly after stimulus onset (see 100ms in Fig. 1A, see Table S1) followed by inferior frontal cortices (Fig. 1, A and C). Prior to speech onset, activity arises in inferior frontal gyrus (IFG) followed by sensorimotor cortices (pre- and post-central gyri; Fig. 1, B and D). During articulation there is a marked suppression of auditory activity (compared with perception), a commonly reported consequence of a corollary discharge (Fig. 1, B to D, and Fig. S1). While these results raise potential candidates for a corollary discharge before speech production across frontal cortices (e.g. IFG, MFG, sensorimotor cortex), local neural activity does not elucidate the exact source and dynamics of the corollary discharge signal.

**Figure 1:**
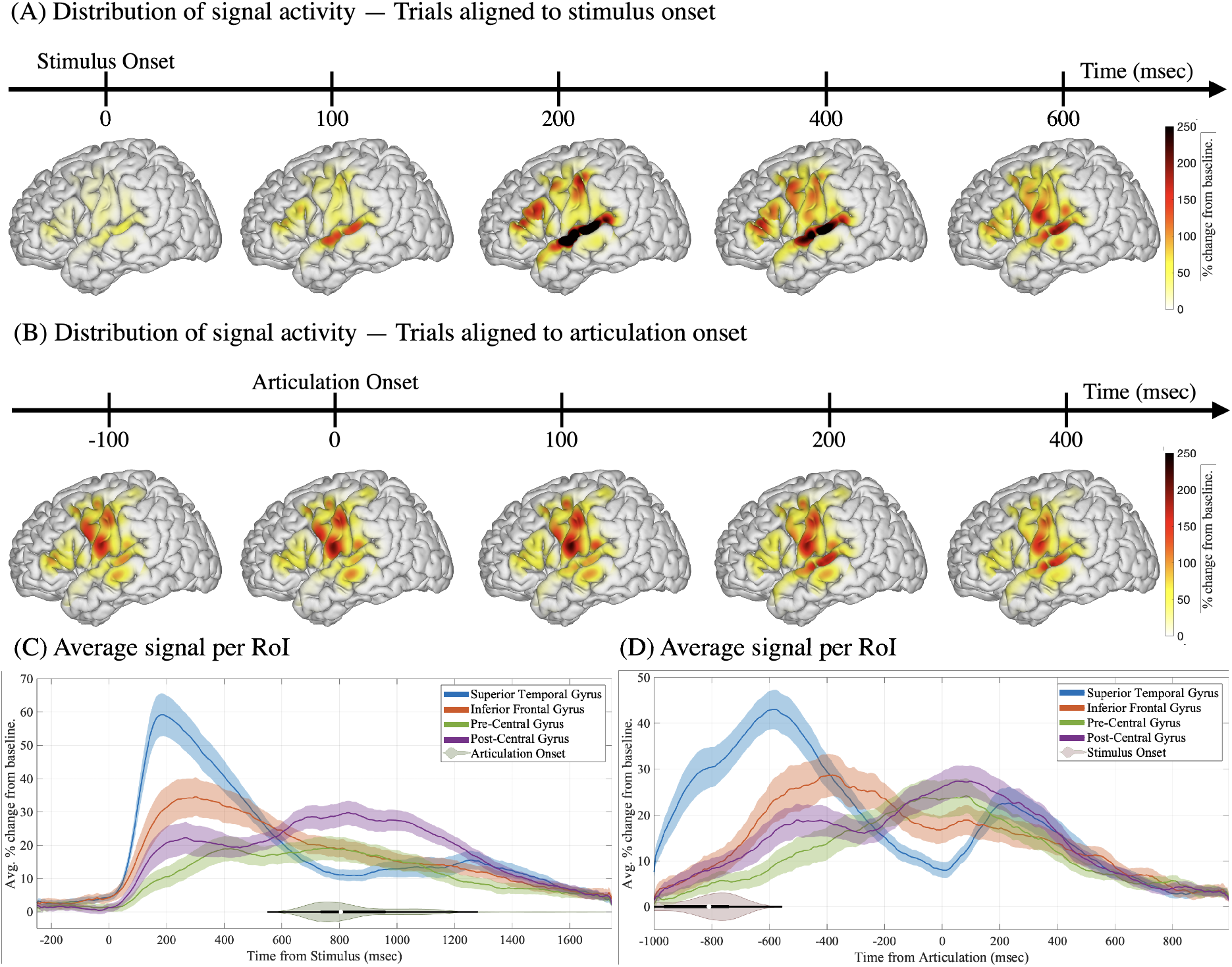
Spatiotemporal distribution of neural activity during speech perception and production. The spatiotemporal distribution of neural activity compared to baseline when participants (N=8) listen to auditory stimuli (A) and subsequently repeat the same word (B). The color code represents percent change from pre-stimulus baseline. A region of interest approach averaging activity in superior temporal, inferior frontal, pre-central, and post-central gyri are shown when trials are aligned to (C) stimulus onset and (D) articulation onset. Shaded regions around the curves depict standard error of the mean across participants. Participants were instructed to repeat the auditory word freely when ready and the reaction time distribution is shown in a horizontal violin plot (C). The stimulus onset relative to articulation onset is shown in the horizontal violin plot in (D).

The corollary discharge signal, by nature, is a blueprint of the motor commands sent to auditory cortex. Hence, a technique is necessary which can measure both the degree of communication between brain regions as well as the direction of information flow. To this end, we developed a directed connectivity analysis framework based on Granger-causality that describes the causal predictive relationships between signals across different electrodes. Unlike previous approaches (e.g. directed transfer function (*23*), partial directed coherence (*24*)) we distilled the large neural connectivity patterns into the dominant communication patterns using an unsupervised clustering technique (orthogonal non-negative matrix factorization (*25*)). Our analysis framework summarizes the dynamics of directed connectivity across cortical regions using a set of connectivity temporal prototypes with corresponding clustering assignment weights (see Fig. 2 and section 4.5 as well as Fig. S2 for methodological details). These prototypes depict typical temporal variation patterns of directed connectivity and their corresponding assignment weights show cortical sources and targets.

**Figure 2:**
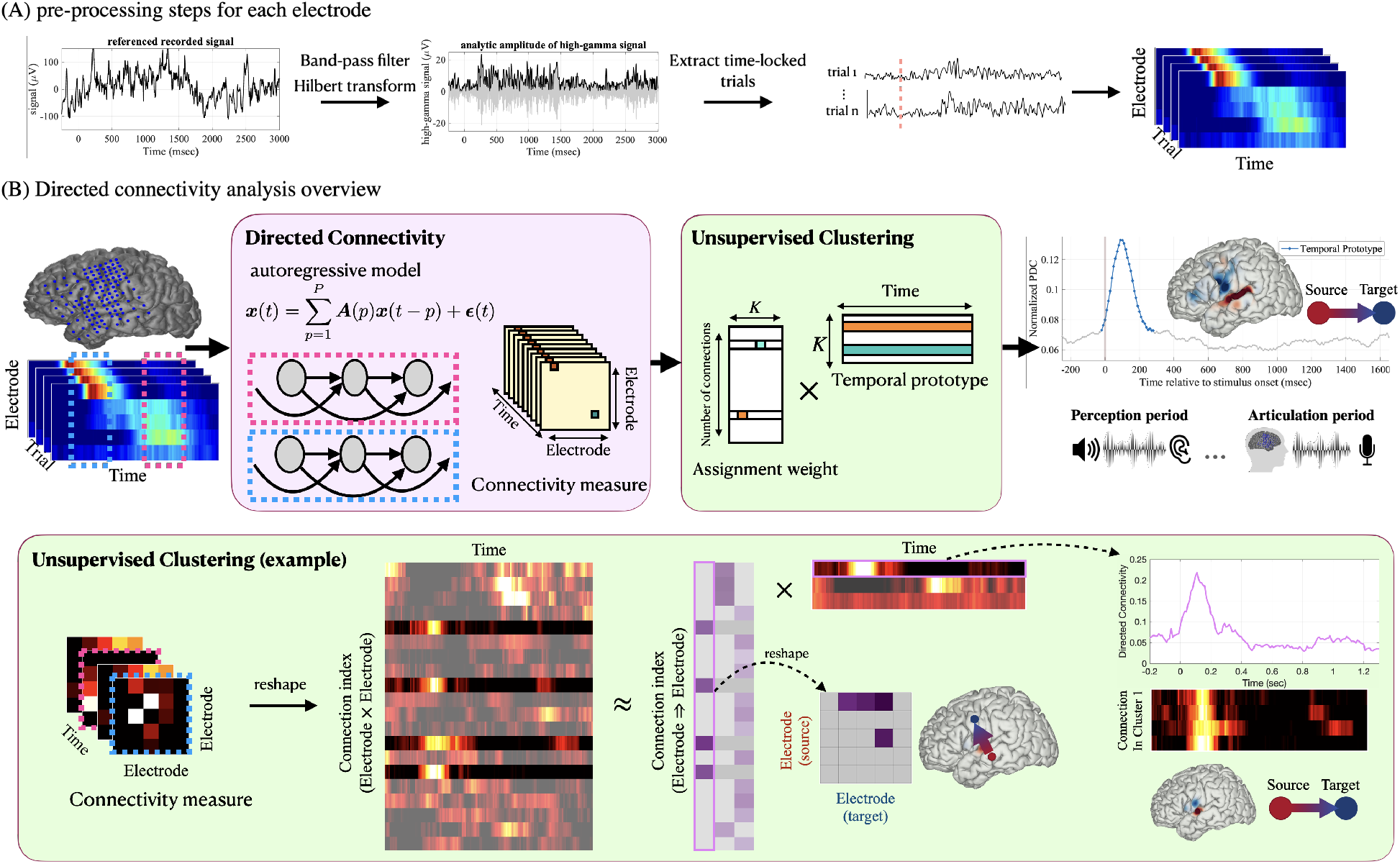
Overview of the signal processing and directed connectivity analysis framework. (A) Electrocorticographic signals are recorded and high-gamma analytic amplitude signal (70-150 Hz) is extracted via band-pass filtering and Hilbert transformation. The signal for each electrode and trial is then extracted locked to perception or articulation onsets. (B) Directed connectivity (based on Granger-causality) is measured using an autoregressive model for successive overlapping time periods, providing the connectivity between different electrodes as a function of time (3D matrix in the purple box). The connectivity patterns are then represented by a few temporal prototypes via an unsupervised clustering technique (orthogonal non-negative matrix factorization; green box). This process reveals the major temporal connectivity patterns via prototypes and their corresponding assignment weights projected on cortex (cortical sources in red and targets in blue). An example of the unsupervised clustering algorithm with connectivity computed for five electrodes is shown. Unsupervised clustering summarizes the connectivity temporal profiles into a few temporal prototypes that represent the temporal changes of those connections. The corresponding assignment weights show cortical sources and targets (connections sourcing from an electrode targeting another).

We first employed our approach in a representative participant (spatiotemporal high gamma activity shown in Fig. 3, A and B), revealing three major connectivity prototypes locked to auditory stimulus onset (Fig. 3C). The first prototype (blue, prototype I Fig. 3C), peaking at 120 msec, showed information flow from STG onto IFG as well as speech motor cortex. The second prototype (yellow, prototype II Fig. 3C), peaking at 340 msec, showed information flow from STG and IFG onto speech motor cortex. The third prototype (purple, prototype III^*†*^ Fig. 3C), peaking at 690 msec, showed information flow from speech motor cortex onto STG. Unlike the high-gamma activity patterns (Fig. 3A), the directed connectivity analysis reveals the source, target, and temporal dynamics of statistically significant information flow (permutation test, *p <* 0.05, see method section 4.5) across cortical regions. Prototypes I and II peak during early auditory processing (while still hearing the auditory stimulus) and exhibit sources from auditory cortex (i.e. STG) implicating these components in auditory comprehension. However, the third prototype shows information flow from speech motor cortex onto auditory cortex and peaks before mean articulation onset (Fig. 3C horizontal violin plot). Neural information flow from motor cortex onto auditory cortex before articulation onset is consistent with the timing and directionality of a corollary discharge signal, providing the first provisional evidence for such a discharge in human speech.

**Figure 3:**
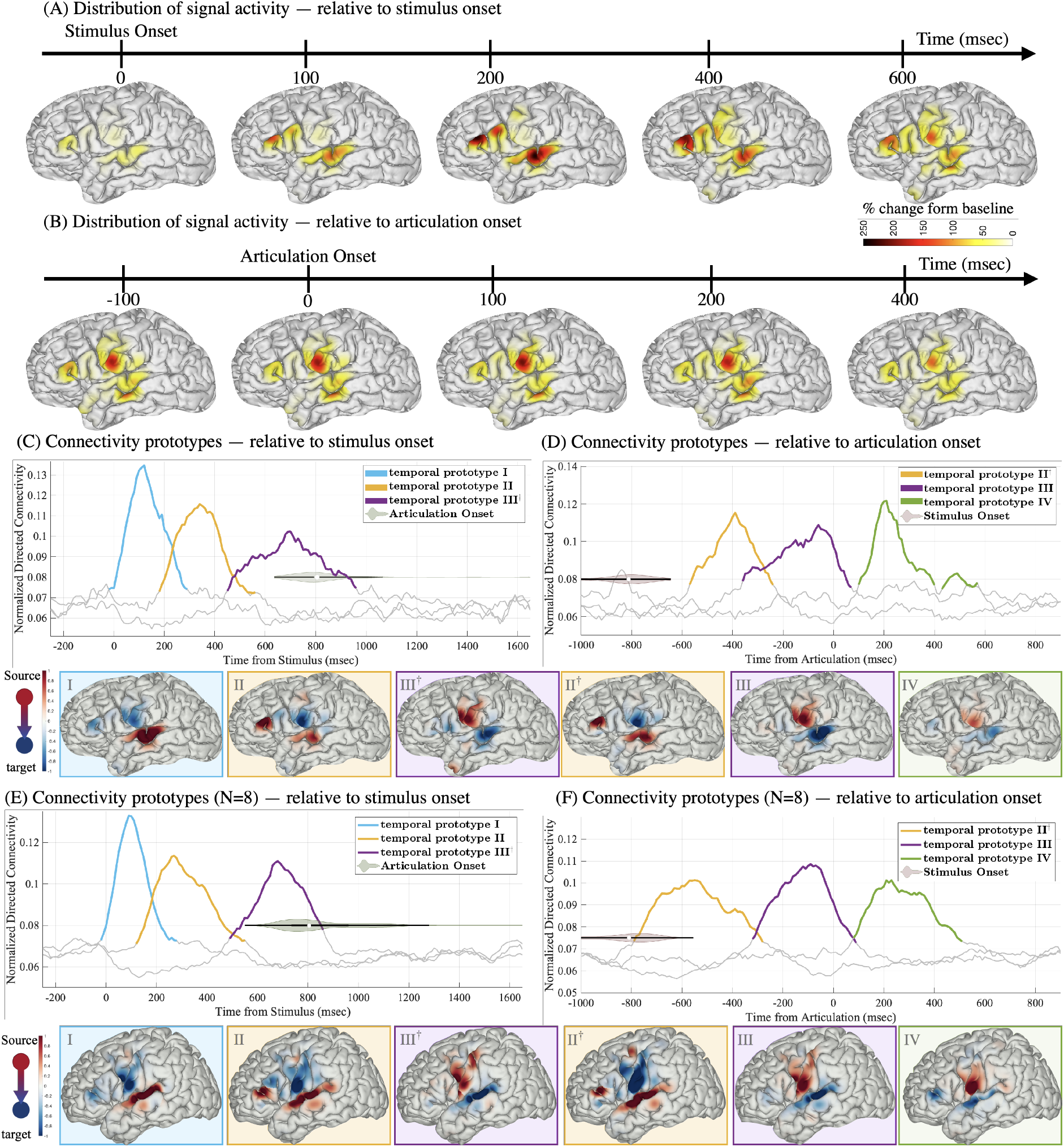
Directed connectivity reveals sources and targets of neural communication during speech perception and production. The spatiotemporal distribution of neural activity compared to baseline in a representative participant is shown during (A) listening to auditory word stimuli and (B) subsequent word repetition. Results of directed connectivity modeling in this participant (temporal prototypes plotted as temporal curves and corresponding information sources in red and targets in blue plotted on cortical surface) are shown locked to (C) stimulus and (D) articulation onset. Colored segments of the curves indicate the portions of the temporal curves that are statistically signifi-cant compared to random permutation (*p <* 0.05, see method section 4.5 for details). (E,F) Directed connectivity analysis repeated for eight participants shows similar prototypical patterns and information sources and targets (vi-sualized on a normal brain). Prototypes I and II (in C and E) show distinct information flow during comprehension part of the task. Prototype II^†^ (in D and F) shows similar timing and information flow related to comprehension and is colored similar to prototype II. Prototype III (in D and F; similar to III^†^ in C and E) is related to pre-articulation, while prototype IV (in D and F) is associated with speech production.

To ascertain the exact timing of information flow related to speech production, we repeated our analysis framework locked to articulation onset. The first connectivity prototype replicated our previous analysis matching information flow from STG and IFG onto speech motor cortex during auditory comprehension (i.e. prototypes II and II^*†*^ in Fig. 3C and D). The next prototype (purple, prototype III Fig. 3D) shows information flow from speech motor cortex onto auditory cortex peaking at -70 msec prior to articulation onset. Lastly, a third prototype (green, prototype IV Fig. 3D) peaking at 210 msec shows information flow from motor cortex onto temporal and inferior frontal areas during speech articulation. Our directed connectivity analysis locked to stimulus and articulation onset reveal multiple prototypes from stimulus onset through production. These prototypes include two related to auditory comprehension (I and II), a pre-articulatory prototype (III) and a speech production prototype (IV). We then replicated this finding across eight participants (clustering the directed connectivity from all the participants together, see Fig. 3, E and F, and Fig. S3 as well as Fig. S4 for variability across participants). To further control for task effects, three of the eight participants performed a passive listening version of the task providing a replication of the two prototypes associated with comprehension (Fig. S5 E, blue and yellow) in overall timing and spatial distribution. Importantly, the passive listening data did not reveal a pre-articulatory prototype (see Fig. S5). The timing before articulation and directionality from motor to auditory cortex of the pre-articulatory prototype (purple, Fig. 3F) establishes a corollary discharge signal replicated across participants.

To verify that the uncovered corollary discharge prototype is not specific to an auditory repetition task, we leveraged a battery of speech production tasks performed by the same participants. These tasks were designed to elicit the same set of matched words during articulation while using various stimulus modalities and word retrieval routes. Participants were instructed to name visual images, read written words, complete auditory sentences, and name auditory descriptions. By clustering the directed connectivity measures across participants for each task, we replicated the corollary discharge temporal prototype and corresponding information flow across all tasks (projected on a template brain shown in Fig. 4, A to E). To investigate how the source varied across sensorimotor cortex and tasks, we analyzed the variance of the corollary discharge’s outflow weights. The source did not differ statistically across tasks (ANOVA main effect of task F(4,369)=1.02, p=0.397). However, we found a significant main effect of region indicating a difference in weight distribution across sensorimotor cortex (dorsal and ventral divisions of precentral and postcentral gyri; ANOVA main effect of region F(3,369)=12.48, p=9.04e-8; no significant interaction F=0.53, p=0.896). The majority of outflow weights originated in ventral precentral gyrus (see Fig. S6 for a post-hoc analysis). To verify the robustness of the corollary discharge temporal profile across participants we clustered the directed connectivity temporal patterns for each participant and task separately. Across participants, we found similar peak timing relative to articulation onset which was not statistically significant across tasks (Kruskal-Wallis test *χ*^2^=6.86, p=0.14), providing an overall mean estimate of, *µ* = -107.5 msec, and peak directed connectivity value, *µ* = 0.1049 (Fig. 4, G and H). Together, these results provide strong evidence across participants and retrieval routes for a corollary discharge signal peaking prior to articulation (−107.5 msec) with neural communication from ventral speech motor cortex onto auditory cortex (Fig. 4F).

**Figure 4:**
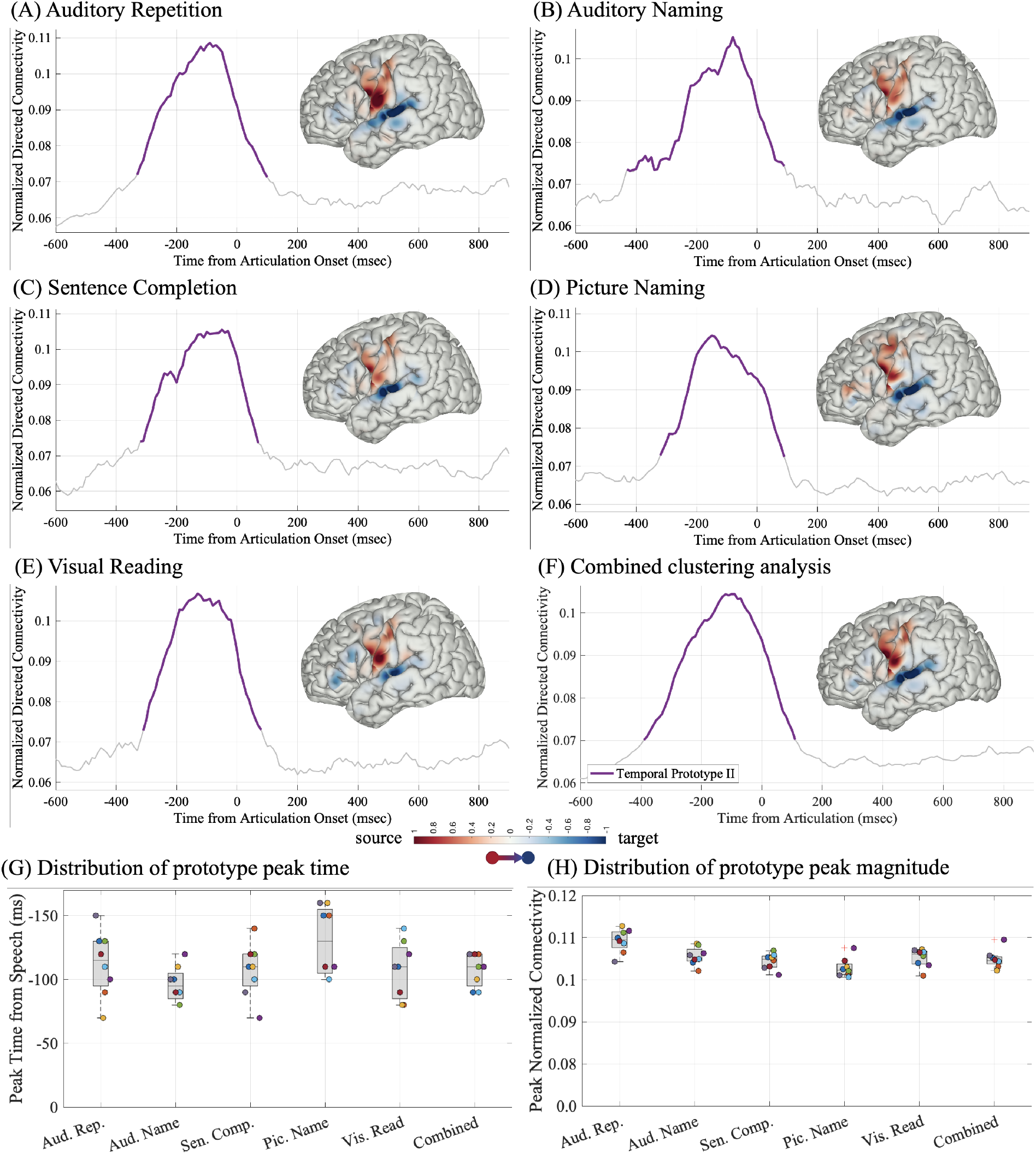
Corollary discharge prototypes are replicated across tasks. The corollary discharge temporal prototype and its corresponding information source (red) and target (blue) are shown for all participants across five different speech production tasks (clustered separately) locked to articulation onset: (A) auditory repetition, (B) auditory naming, (C) sentence completion, (D) picture naming, and (E) visual word reading. (F) An overall representation of the corollary discharge prototype is shown by clustering the combined connectivity results from all tasks and participants. Colored regions of the curve represent statistically significant directed connectivity compared to random permutation (*p <* 0.05). The distribution across participants of the peak time (G) and magnitude (H) of the corollary discharge prototype is shown for each task (clustered for each participant separately; circles are color-coded per participant; Box-plots show minimum, maximum, quadrants, and median). The combined peak time and magnitude are established by clustering within participant but combining all the task data for the last column of G, H.

To date, there was a lack of direct evidence for a corollary discharge signal and its exact location, although its consequence – a suppression of auditory cortex during speech production – has been reported (*4, 14, 26*). To verify that the novel prototype we discovered is the source of a corollary discharge we sought to show a direct link to speech induced auditory suppression. Previous human (*14*) and non-human primates (*12*) studies reported a wide range of suppressed responses in auditory cortex. We hypothesized that the degree of information flow from the corollary discharge source should predict the intensity of suppression across auditory cortex recording sites. We first quantified the level of neural activity for each electrode anatomically located within auditory cortex (i.e. within participant parcellation of superior temporal gyrus, see Fig. 5, A and B) and quantified the level of suppression by computing a normalized index (varying between 1 for completely suppressed and -1 for completely enhanced, see Fig. 5, C and D). We then quantified the degree of information flow into each electrode in auditory cortex (i.e. inflow connectivity weights see Fig. 5E; see methods 4.6) from the sources in our corollary discharge prototype (temporal prototype III in Fig. 3F). To test our hypothesis, we correlated the degree of information flow into each auditory electrode with its level of suppression and found a significant relationship (Pearson correlation coefficient r=0.430, p=1.45e-4, see Fig. 5F; see methods 4.6). This further establishes that information flow from speech motor cortex before articulation promotes auditory suppression during speech. A majority of the outflow over speech motor cortex originated in ventral motor cortex (Fig. 5G). To verify the timing of this suppression signal, we repeated the correlation analysis across multiple time windows while ensuring that the information flow originated from ventral motor cortex and found significant correlations (Permutation test, p*<* 0.05; see methods 4.6) peaking before speech articulation (Fig. 5H).

**Figure 5:**
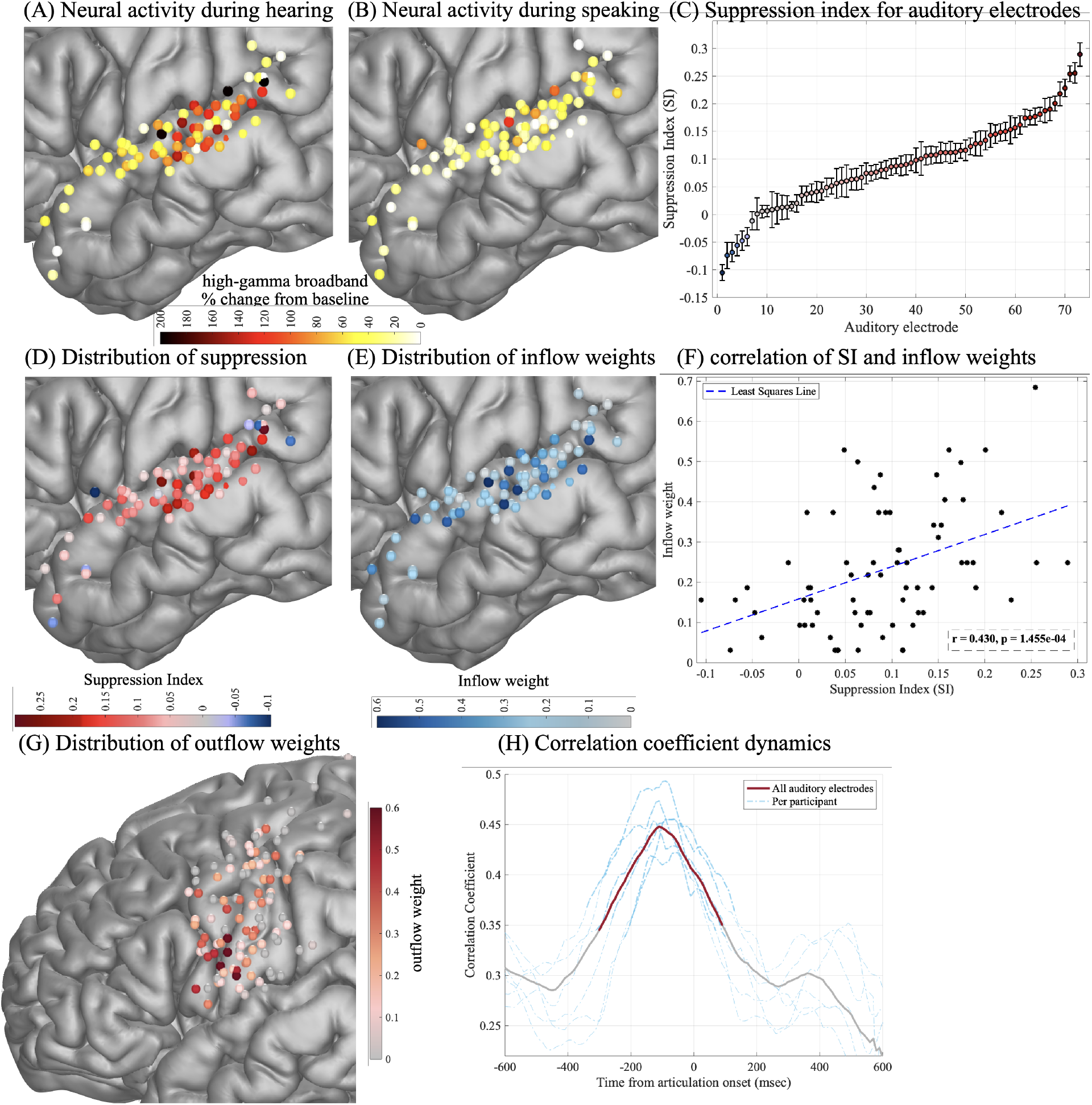
Corollary discharge predicts speech-induced auditory suppression. Average neural activity when participants (A) hear auditory stimuli and (B) speak during an auditory repetition task. (C) Most auditory responses are suppressed during speech production, quantified by a normalized suppression index (SI) shown for all active auditory electrodes (i.e. within participant anatomical label of Superior Temporal Gyrus) across all participants. Electrodes are sorted according to their mean SI over trials (error-bars represent SEM across trials) and vary from 1 (completely suppressed) to -1 (completely enhanced). Spatial distribution of mean SI for auditory electrodes across participants (D) and of the inflow weights onto auditory electrodes (E) in the corollary discharge prototype III (purple curve in Fig. 3F). (F) The correlation between mean SI and the inflow weights of the corollary discharge for all auditory electrodes. (G) Spatial distribution of the outflow weights (i.e. source) of the corollary discharge prototype in pre- and post-central gyri. (H) The dynamics of the correlation coefficient between mean SI and directed connectivity from sources in ventral motor cortex onto auditory cortex as a function of time (solid curve is obtained using all electrodes across participants and dashed lines show the analysis per participant; red shading denotes statistical significance from a permutation test *p <* 0.05).

## 3 Discussion

Corollary discharge signals from motor onto auditory neural populations are a hallmark of neural circuitry in animals and convey information on impending actions (*4, 27, 28*). In humans however, the exact source and dynamics of the signal remained unknown. We leveraged the excellent spatiotemporal resolution of electrocorticography recordings from neurosurgical patients and developed a novel signal analysis framework to study the dynamics of corollary discharge signal in human speech. Our analysis of neural connectivity dynamics revealed four distinct stages during auditory repetition likely representing comprehension (two stages), pre-articulatory preparation, and speech production per se. A cardinal connectivity temporal prototype during the pre-articulatory stage showed information flow from speech motor cortex onto auditory cortex before articulation onset. This prototype was replicated across participants and speech production tasks even when varying stimulus modality and routes of word retrieval (i.e. repetition, naming, completion), highlighting ventral pre-central gyrus as the source of the corollary discharge. Lastly, we directly linked the information flowing from this discharge with the degree of auditory cortical suppression in each electrode. Together, these findings depict the timing, source, and consequence of the corollary discharge signal in human speech.

We designed a novel directed connectivity analysis framework to study the cortical information flow by fusing techniques that analyze the causal relationships between time-series based on Granger-causality measures (*23, 29–34*) and unsupervised clustering (orthogonal non-negative matrix factorization (*25*)). Only a select few previous ECoG publications have leveraged similar effective connectivity between brain regions during language tasks (*35, 36*), revealing the role of Broca’s area in coordinating articulately preparation (*22*) as well as frontal orthographic processing during word reading (*37*). In addition to replicating previous findings, we are able to do so on the single participant level. In addition to previous findings of early temporal to inferior frontal flow (*22*) we find additional temporal to Rolandic cortex information flow, albeit not causing downstream suppression. A major limitation of previous connectivity approaches is the large set of revealed connections rendering the results difficult to interpret and test statistically without averaging across cortical regions. Our unsupervised clustering approach circumvents this issue by finding a select few source-and-target pairs of information flow representing the prototypical temporal dynamics of connectivity across cortex. Our approach identified cortical communication supporting speech preparation (*22*), within a single and across participants. Further, it revealed a novel corollary discharge signal, possibly obscured in the past due to limited cortical coverage as well as averaging across regions and participants.

Theoretical models of speech production all assume a corollary discharge or a motor initiation signal, presumably originating from frontal cortex and targeting the auditory sensory areas (*38–40*). However, the exact source and timing dynamics of this signal is variable and unclear due to the lack of direct evidence in neural recordings. Evidence for corollary discharge has been limited to two major results in both human and non-human primates: suppression of auditory cortex during vocalization (*4, 13–15, 41–45*), and enhanced auditory responses during altered feedback (*7, 8, 12, 46, 47*). Our results showed a distinct source of information flow originating in ventral pre-central gyrus before articulation irrespective of task modality. While we cannot rule out sub-cortical sources that were not recorded, we did not find other sources across cortical peri-sylvian sites. Specifically, inferior frontal gyrus showed major information flow during the comprehension prototypes but critically, not during the pre-articulatory prototype. While IFG has been implicated as a major cortical source of connectivity during stimulus comprehension and pre-articulation (*22, 37*), our data would rule out its involvement as a corollary discharge source. This is further supported by our consistent corollary discharge prototype across tasks, in face of previous reports of attenuated activity during visual tasks (*37, 48*).

Further, our reported location and timing are consistent with the hypothesized communication across theoretical frameworks, as well as neuronal suppression which has been reported prior to vocalization (*4*). In stark contrast to prior studies which failed to link motor neural responses with auditory suppression (*8, 14, 16*), our results clearly establish that the target of information flow correlates with the degree of suppression. This correlation peaks during the pre-articulatory period and levels out during speech production. While this link is a critical litmus test for corollary discharge, the correlation timing suggests that a local circuit mechanism may sustain suppression during production.

Neurons supporting a corollary discharge circuit have been established in the cricket (*2*), songbird (*49*), and several other mammals (*1*). In humans, a corollary discharge signal is involved in speech production as well as the ability to distinguish between self- and external-generated thoughts and actions (*10, 50*). A dysfunctional corollary discharge circuit is the major model explaining auditory hallucinations in schizophrenia patients (*10, 51–55*). Auditory suppression is impaired in schizophrenia (*10, 11, 52*), but no direct corollary discharge signal source has been identified. Our novel framework and results provide a missing link in the human auditory system and elucidate the source, target and timing of the corollary discharge network with major implications for dysfunction of speech and psychosis.

## 4 Methods

### 4.1 Participant information

A total of 8 neurosurgical patients (7 female, all with left hemisphere coverage, mean age: 38 with range 19 to 55 years) were implanted with electrocorticography electrodes and provided informed consent to participate in this research. All consent was obtained in writing and then requested again orally prior to the beginning of the experiment. Electrode implantation and location were guided solely by clinical requirements. Five of the participants were implanted with standard clinical electrode grid with 10 mm spaced electrodes (Ad-Tech Medical Instrument, Racine, WI). Three participants consented to a research hybrid grid implant (PMT corporation, Chanassen, MN) that included 64 additional electrodes between the standard clinical contacts (with overall 10 mm spacing and interspersed 5 mm spaced electrodes over select regions). This provided denser sampling of underlying cortex but was positioned solely based on clinical needs. The superior temporal gyrus (STG) region is sampled for all participants, and other cortical regions (including Broca’s area and motor cortex) are also sampled. The study protocol was approved by the NYU Langone Medical Center Committee on Human Research.

### 4.2 Experiment setup

The participants were instructed to complete five tasks to pronounce the target words in response to certain auditory or visual stimuli. These five tasks consisted of: auditory repetition (i.e., repeat an auditory presented word), auditory naming (i.e., naming a word based on an auditory presented description), sentence completion (i.e., complete the last word of an auditory presented sentence), visual word reading (i.e., read out loud a visually presented written words), and picture naming (i.e., name a word based on a visually presented line drawing). All tasks consisted of the same 50 target words. Each trial began with the stimulus presentation and participants were instructed to respond freely when they were ready. For each task and trial the stimulus was randomly presented. Each target item appeared twice in the auditory repetition, picture naming, and visual word reading tasks and once in auditory naming and sentence completion.

### 4.3 Data collection and general pre-processing

The ECoG recordings were collected as the participant performed each task. As an initial inspection, we rejected electrodes with epileptiform activity, line-noise artifacts, poor contact, and high amplitude-shifts. These initial criteria were used based on the input from the clinical team to remove electrodes that are visually and obviously problematic and outliers. The exclusion of electrodes with epileptiform activity was done based on the characterization provided by the clinical team as interictal population or seizure onset zone. ECoG recordings were referenced using the common average reference approach by averaging the signal across all electrodes and subtracted from each individual electrode signal. The raw voltage signal from each electrode was then filtered to high gamma broadband range (70-150 Hz) and the analytic amplitude (envelope) signal was then extracted by a Hilbert transform. The envelope signal was then downsampled to 200 Hz (see Fig. S7 for an exemplar electrode). The high-gamma analytic amplitude (envelope signal) has a band limit of 80 Hz, so down-sampling to 200 Hz (above the Nyquist rate) after amplitude extraction does not affect the frequency content (see Fig. S7 C). This down-sampling procedure reduces computational complexity and enhances the numerical stability of the autoregressive models (*35, 36*). The continuous data stream was divided into epochs based on the onset of stimulus (locked to stimulus) or onset of speech (locked to articulation). We restricted our analysis to a subset of active electrodes that showed strong event-related activity when averaged over trials (see supplemental text B for detailed explanation of the selection criteria as well as Fig. S8 and Fig. S9 for examples of active electrodes and the overall spatial distribution).

### 4.4 Neural activity visualization

When plotting the neural activity (mainly shown in Fig. 1, Fig. 3 A-B, and Fig. 5 A-B), we presented the data in units of percent change from baseline activity. We performed the general pre-processing steps introduced in section 4.3. We then normalized each trial by the mean activity in that trial’s baseline (250 msec before stimulus presentation). In Fig. 1, A and B, this normalized signal (presented in units of percent change from baseline) for each electrode is averaged over trials and time in a 50 msec window centered at each marked time-stamp and projected onto a normal brain with a Gaussian kernel of size 50 mm (we restrict the spread of the Gaussian kernel to within the boundaries of the associated cortical region for each electrode). Similarly, when showing a representative participant in Fig. 3, A and B, the same procedure is performed for the electrodes and Cortical surface model of the participant’s brain. In Fig. 1, C and D, the normalized broadband high-gamma envelope signal (in units of percent change from baseline) is averaged over electrodes within a region of interest and trials for each time-point and participant. We show mean and standard error of the mean across participants for each region of interest. The distribution of articulation onset relative to stimulus onset (Fig. 1C, Fig. 3C and E) and stimulus onset relative to articulation onset (Fig. 1D, Fig. 3 D and F) over trials and participants are shown as horizontal violin plots.

### 4.5 Directed connectivity analysis framework

Here, we first provide a brief overview of the steps used for directed connectivity analysis framework. We then provide detailed explanation of each step in the following subsections.

For directed connectivity analysis, we performed the general pre-processing steps introduced in section 4.3. We z-scored the signal from each electrode by the mean and standard deviation from all the time-points and trials. For each participant and task, we used the trial information to fit a multivariate autoregressive (MVAR) model to 100 msec overlapping time windows with hops of 10 msec (see section 4.5.1 for details of model fit). For each window we measured the directed connectivity (Granger-causal sense) between electrodes by computing the partial directed coherence (PDC) (*24*) from the fitted MVAR model coefficients (see section 4.5.2 for details). We focused our analysis on temporal changes of PDC as the resulting connectivity showed the information flow between any two given electrodes as a function of time. Our goal was to summarize the data into a few prototypes that represent the major temporal changes of connectivity. To derive the prototypical temporal connectivity patterns, we used orthogonal non-negative matrix factorization (ONMF) (*25*) as an unsupervised clustering technique. Similar to other dimentionality reduction algorithms such as principal component analysis (PCA), ONMF summarized the temporal connectivity patterns. However, a distinct feature of ONMF, in contrast to PCA, is that each connection is associated with one and only one prototype resulting in a clustering algorithm. We gathered the measured directed connectivity signals into a matrix (the dimensions of which are number of time-windows by number of connections) and applied the ONMF, representing each connection as a scaled version of one of a prototypical patterns (each connection can only be assigned to one prototype; see section 4.5.3 for details). Consequently, ONMF clustered the temporal PDC changes into a few clusters. For each cluster, a temporal prototype represents the temporal behavior (“when”, *K × T* matrix in Fig. S10) of that cluster, and assignment weights show which connections belong to that cluster (“where”, *M* (*M* − 1) *× K* in Fig. S10). The assignment weights associated with each cluster are then visualized on participant or Montreal Neurological Institute (MNI) brain with a Gaussian kernel of size 50 mm (restricting the spread to within the boundaries of the associated cortical region for each electrode).

#### 4.5.1 Multivariate autoregressive model

We denote by *x*_*m*_(*t*) ∈ *ℝ*_≥0_ the envelope of the high-gamma broadband extracted from the ECoG recordings, downsampled to 200 Hz, and Z-scored based on mean and standard deviation per electrode. The subscript _*m*_ denotes the active electrode index *m* ∈ {1, *…, M* } and *t* the time-step *t* ∈ {1, *…, T* }. The signal *x*_*i*_ Granger causes the signal *x*_*j*_ if knowledge of *x*_*i*_(*τ*) for *τ* ≤ *t* improves the prediction of *x*_*j*_(*t*). To assess Granger causality between multichannel ECoG data, we fitted a multivariate autoregressive (MVAR) model. Let 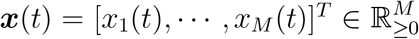 be the multichannel high-gamma analytic amplitude signal at time *t* with *M* total active electrodes (*M* total channels). The MVAR model assumes that the signal at each time point, ***x***(*t*) can be estimated as a linear combination of the signal at previous time-points and a random innovation signal ***ϵ***(*t*). Consequently, we can write the MVAR formulation as

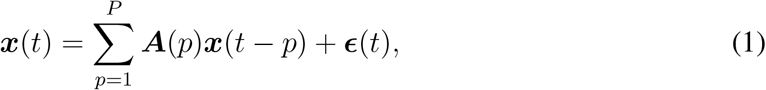

where ***A***(*p*) ∈ ℝ^*M×M*^ are coefficient matrices for which the element *a*_*ij*_(*p*) shows the dependency of *x*_*i*_(*t*) on *x*_*j*_(*t*−*p*) for electrodes *i, j* = 1, *…, M* and time-lags *p* = 1, *…, P*. The random innovation signal ***ϵ***(*t*) is assumed to be composed of white uncorrelated noises with covariance matrix ***Q***. Causality relations are found when the relevant interaction is active in (1). More formally, we can say *x*_*j*_ Granger causes *x*_*i*_ if *a*_*ij*_(*p*)*≠* 0 at least one *p* ∈ {1, *…, P* }. This is consistent with the definition of direct causality as in (*56*). The parameters of the MVAR model in (1), Θ = {***A***(*p*), ***Q***}, are estimated using the Expectation-maximization algorithm (*36, 57*).

The AR process is a linear model with an inherent stationarity assumption on the signal ***x***. We follow the recommendation of Ding et al. to model short windows of signal with separate AR models (*57*). We used short overlapping windows of 100 msec with hops of 10 msec and used the available trial data to fit an MVAR model for each window. This allowed us to look at dynamic changes of connectivity across larger time-scales of the entire trial.

We followed the recommendations in (*35, 58*) for choosing the model order (based on AIC/BIC criterion). Specifically, for each 100 msec window (non-overlapping windows were used for model order selection), we separately computed the model order that minimizes AIC and BIC criteria (tested over range 1 to 10) and picked the median across windows. In the event that the median from AIC and BIC did not match, we chose the overall median. We typically found the model order of *P* = 4 corresponding to delays up to 20 msec. We check the stability of the estimated MVAR model for each window by computing the roots of the characteristic polynomial

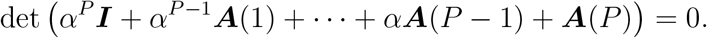

We made sure that the roots satisfied |*α*| *<* 1 for each window. In the very rare event that this condition is not satisfied the model order is decreased and the model fit repeated. We performed the Ljung-Box portmanteau test for whiteness and the Kendall’s *τ* test for independence (*59*) on the resulting MVAR model residual ***ϵ***(*t*) to ensure temporally uncorrelated residual.

#### 4.5.2 Measuring directed connectivity from model coefficients

We used partial directed coherence (PDC), a frequency-domain approach to describing the relationships (direction of information flow) between multivariate time series (*24*), to measure directed connectivity. We computed the PDC from the fitted MVAR coefficients of each window as

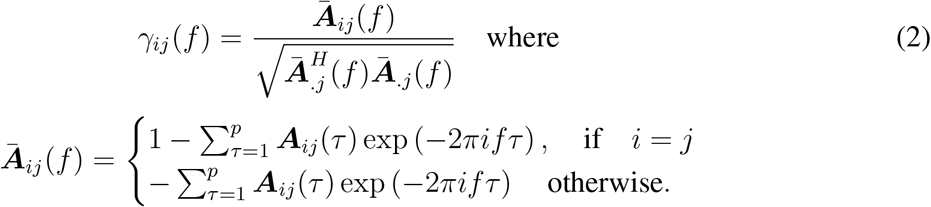

To obtain a picture of how the connectivity changes across time-windows, we followed (*36*) and defined *γ*_*ij*_ = Σ_*f*_ |*γ*_*ij*_(*f*)| (summing the PDC values over frequency for each window) and focused on the temporal changes of *γ*_*ij*_ over the shifted windows. We note that PDC satisfies 0 ≤ |*γ*_*ij*_(*f*)|_2_ ≤ 1 and 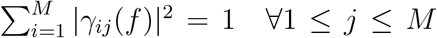 (normalized interaction strengths with respect to a given signal source).

#### 4.5.3 Clustering the temporal patterns of directed connectivity

We were interested in changes of PDC values as a function of time. We used unsupervised clustering, performed by orthogonal non-negative matrix factorization (ONMF), to find and group connections with similar temporal patterns of PDC changes into one cluster. For each time window *t* ∈ {1, *…, T* }, let 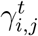 be the PDC computed from node *i* ∈ {1, *…, M* } to node *j* ∈ {1, *…, M* }. We formed the matrix 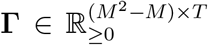 such that each row represents the temporal changes of PDC between a particular pair of electrodes; i.e. 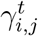 for the connection from node *i* to node *j* with *i≠ j*. To cluster similar temporal PDC profiles (i.e. similar rows in **Γ**) into the same group, we used orthogonal non-negative matrix factorization (ONMF). For a desired number of clusters *K*, the ONMF objective function can be written as,

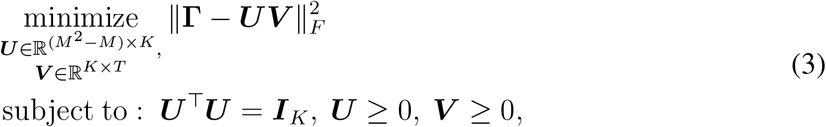

where the constraints ***U*** ≥ 0 and ***U*** ^⊤^***U*** = ***I***_*k*_ imply that each rows of ***U*** can have at most one non-zero entry, hence the clustering nature of this objective function. Each cluster *k* is represented by its corresponding row ***v***_*k*_ in ***V***, which we refer to as the *k*-th temporal connectivity prototype.

We used the EM-ONMF algorithm (*25*) to solve for the matrices ***U*** and ***V***. This algorithm finds a set of disjoint clusters, *π*_*k*_, *k* = 1, 2, …, *K*, such that each cluster *π*_*k*_ contains rows in **Γ** that are as similar to each other as possible. If the temporal connectivity profile from node *n* to node *m* is clustered to cluster *k*, then it can be approximated by 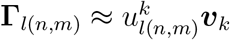. We call the weight 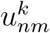 the assignment weight for prototype *k* from *n* to *m*. We define the inflow weight for each electrode *m* in cluster *π*_*k*_ as 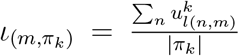, where |*π*_*k*_| indicates the number of connections in cluster *π*_*k*_. Similarly we define the outflow weight for each electrode *n* as 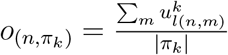. To visualize the source and target locations associated with each prototype, we defuse the inflow (negatively weighted; blue color) and outflow (positively weighted; red color) weight associated with each electrode into the surrounding area in the same anatomical region using a Gaussian kernel on a patient or standard brain map.

We assessed the statistical significance of prototypes with a permutation test (*60*), where we randomized the cluster assignments and recomputed the prototypes for 1000 repetitions. We tested each time-point of the prototype against the randomized projection distribution (with an alpha criterion of 0.05) and to control for multiple comparisons error only continuous range of values (longer than 100 msec, corresponding to 5 windows) showing statistical significance was accepted. The number of clusters (corresponding to total number prototypes) were determined based on the relative reconstruction error 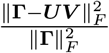 for different number of clusters. We empirically found that this error term plateaued at four components for the time windows considered in our experiments (see Fig. S11). One of the resulting prototypes always showed “noise-like” activity and was removed from analysis (see Fig. S3).

### 4.6 Speech induced auditory suppression

For each electrode anatomically located in the auditory cortex (i.e. STG) we obtain the broadband high-gamma signal by applying the general pre-processing steps in section 4.3. To quantify the level of suppression for each auditory electrode, we compute the average broadband high-gamma signal in 300 msec after the stimulus onset 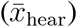 and 300 msec after articulation onset 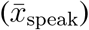 in the auditory repetition task. For each auditory electrode (total of 73 across all participants) we compute the suppression index (SI) defined as 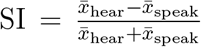 (varying between 1 for completely suppressed and -1 for completely enhanced; distribution shown in Fig. 5, C and D). We quantify the information inflow to each auditory electrode by the cluster assignment weight associated with the corollary discharge prototype from auditory repetition task (distribution shown in Fig. 5E). To assess the link between suppression and corollary discharge we use Pearson correlation across all auditory electrodes for all participants (Fig. 5F). To show the temporal dynamics of this correlation, for each auditory electrode we compute the average directed connectivity of the connections represented by the corollary discharge prototype sourcing from electrodes located anatomically in ventral pre-central gyrus in overlapping windows of 300 msec with hops of 10 msec. We then correlate the suppression index and the average directed connectivity for each time-window across all auditory electrodes (solid curve in Fig. 5H). We repeat the same analysis only considering the auditory electrodes for each participant (dashed curves in Fig. 5H; two participants were removed from this analysis due to limited number of auditory electrodes). We assess the statistical significance by a permutation test (randomizing the suppression index assignment for each window a total of 1000 times and comparing each time point to this distribution; alpha of 0.05; multiple comparisons error corrected with only accepting continuous range longer than 100 msec).

## Acknowledgements

This work is supported by National Science Foundation IIS-1912286 and the National Institute of Health NINDS 1R01NS109367, 1R01NS115929, R01DC018805 grants. The authors would like to thank Dr. David Schneider for his valuable comments on this manuscript.

## Data availability

Data to replicate figures in this manuscript is available at: https://github.com/flinkerlab/AuditoryCorrolaryDischarge.

## Code availability

The code and sample data for the directed connectivity analysis framework is freely available from: https://github.com/flinkerlab/AuditoryCorrolaryDischarge.

## Author contributions

A.K-G. proposed and implemented the directed connectivity analysis framework and the unsupervised clustering algorithm with advisement from Y.W. and A.F.; R.W. and X.C participated in the data processing; L.Y. participated in data acquisition and preprocessing; P.D. and D.F. provided clinical care; W.D. provided neurosurgical clinical care; O.D. assisted in patient care and consent; Y.W. co-led the research project with A.F. and advised from engineering perspective; A.F. led the project, participated in all data acquisition, and advised from neuroscience perspective; A.K-G. and A.F. co-wrote the manuscript with input from all authors.

## Supplementary Materials

Figures S1 to S11

Table S1

## Supplementary Materials for

## S Supplemental Data

**Figure S1:**
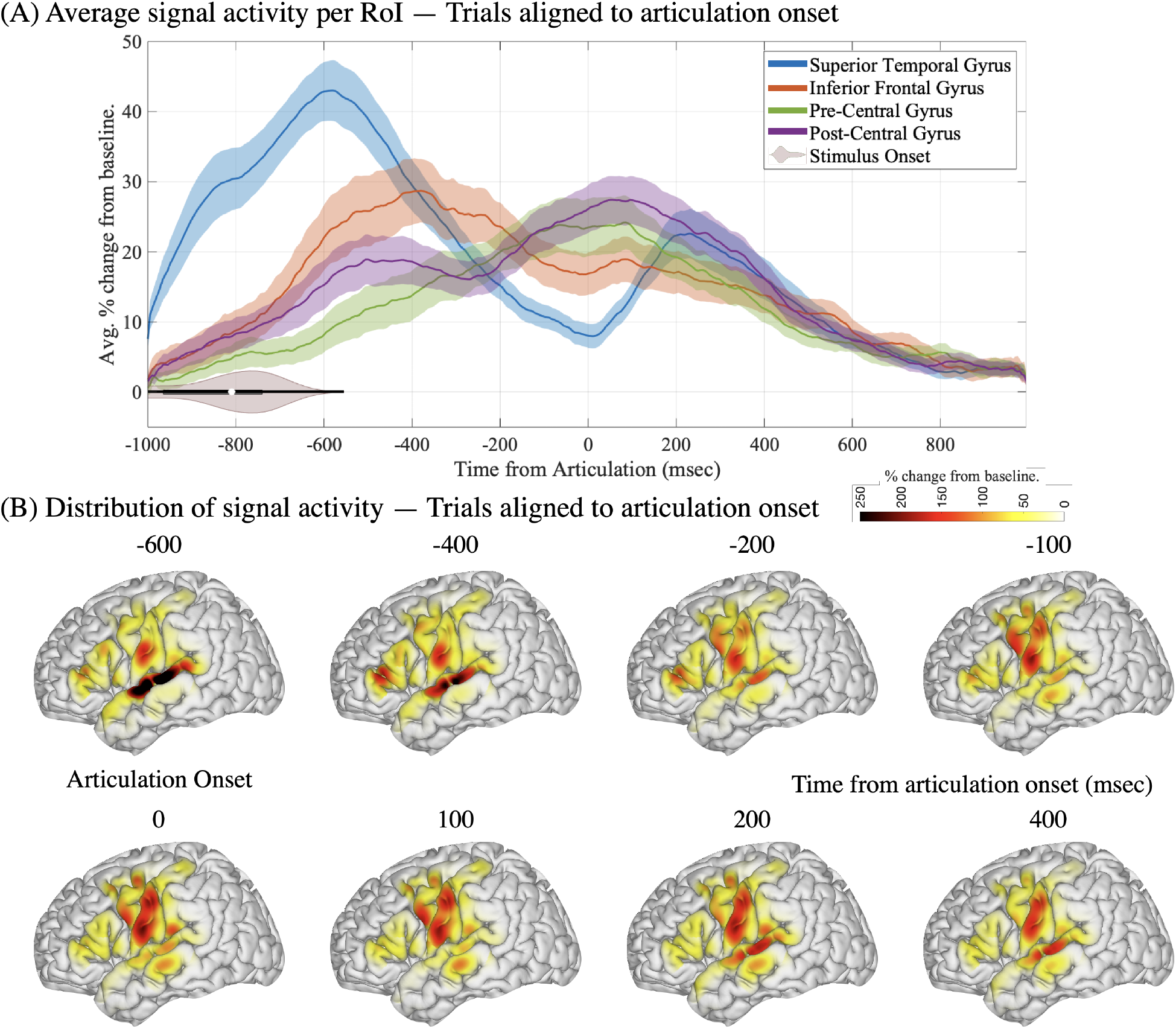
Spatiotemporal distribution of neural activity before and after speech production. A region of interest approach averaging activity in superior temporal, inferior frontal, pre-central, and post-central gyri are shown when trials are aligned to articulation onset (A). The spatiotemporal distribution of neural activity compared to baseline when participants articulate are shown in (B). The color code represents percent change from pre-stimulus baseline.

**Figure S2:**
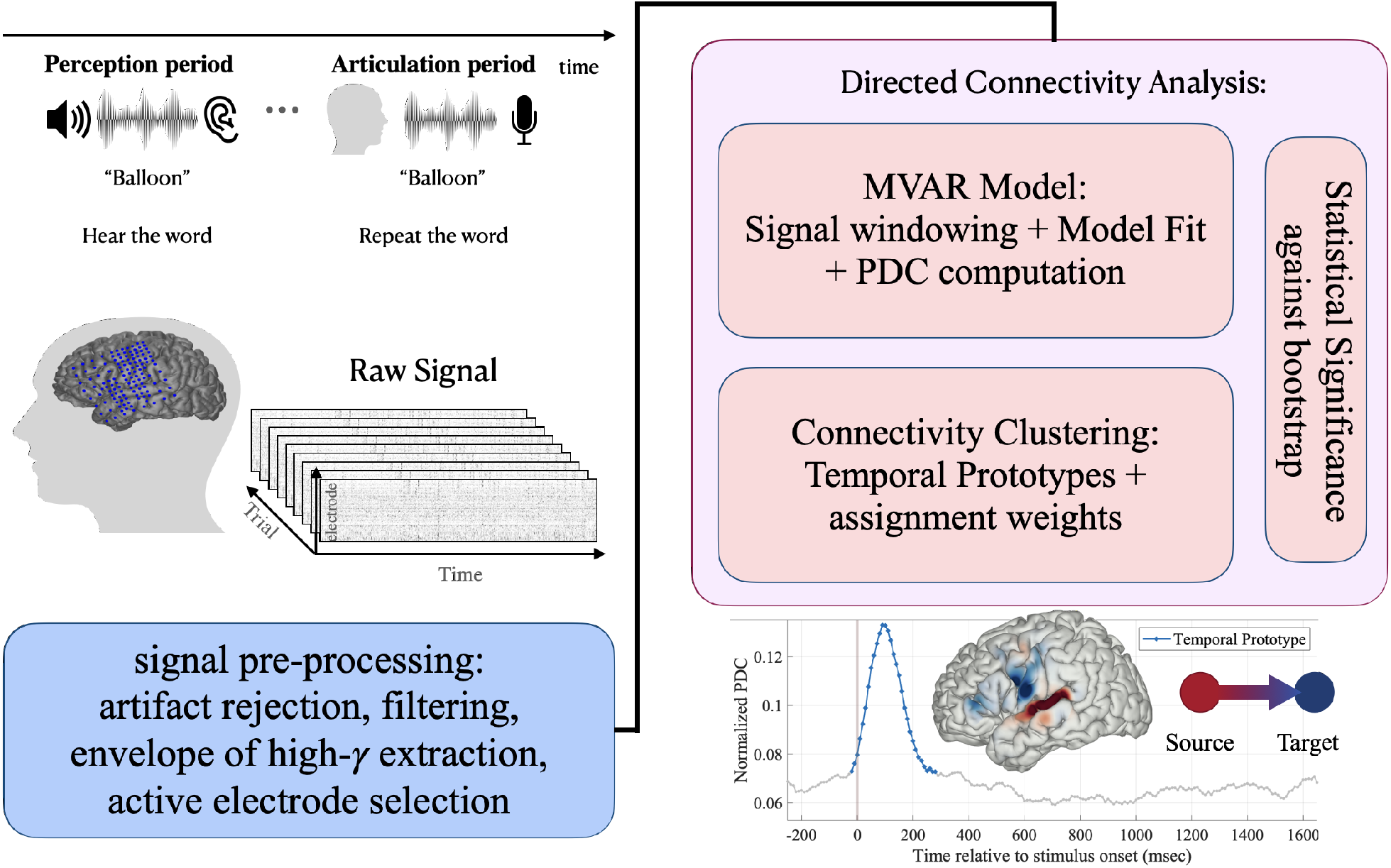
Detailed overview of the proposed directed connectivity analysis framework. Electrocorticographic signals are recorded from neurosurgical patients. We focus on the analytic amplitude of high-gamma broadband locked to perception or production onset. Directed connectivity is measured using an autoregressive model for successive overlapping time windows. The connectivity patterns are then represented by a few temporal prototypes via an unsupervised clustering technique. This process reveals the major temporal connectivity patterns and their corresponding assignment weights projected on cortex. We test statistical significance of each prototype against random permutation (*p <* 0.05).

**Figure S3:**
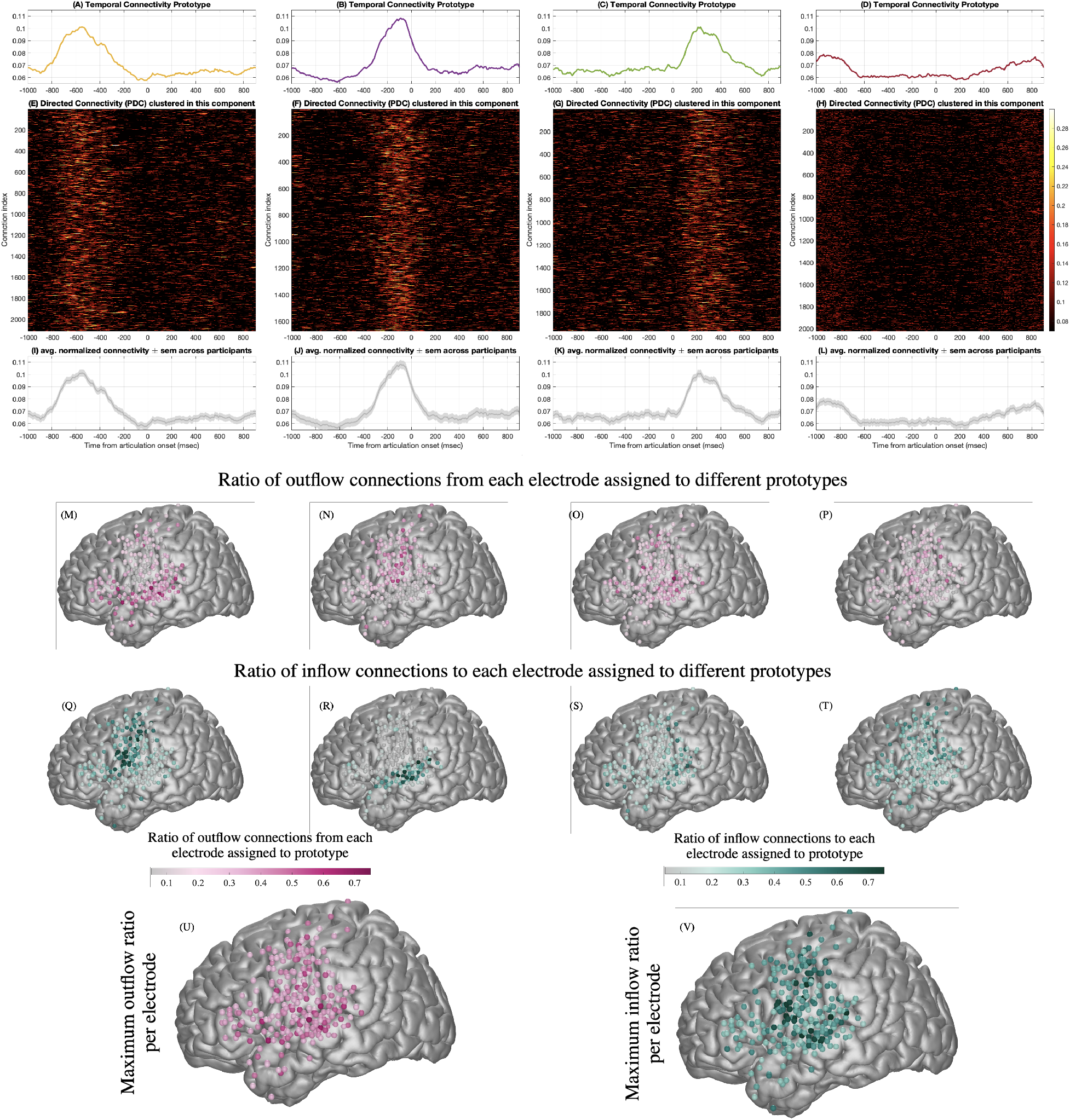
Temporal connectivity profiles in auditory repetition locked to articulation. (A-D) Resulting temporal connectivity prototypes when directed connectivity profiles from auditory repetition task locked to articulation are clustered to 4 clusters for eight participants together. (E-H) The temporal connectivity profiles, 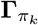, assigned into cluster *k*, represented by the corresponding prototype in A to D plotted as images. Each row in matrices E to H shows the temporal dynamics of a specific connection between two electrodes in each cluster. The rows of the matrices are ordered by participants. (I-L) The average normalized connectivity and corresponding standard error of the mean across participants (shaded region) shows robustness of the corresponding prototypes. (M-P) Ratio count of outflow connections from each individual electrode that were assigned to each corresponding prototype. (Q-T) Ratio count of inflow connections to each individual electrode that were assigned to a prototype. Maximum ratio of outflow (U) and inflow (V) across prototypes for each electrode.

**Figure S4:**
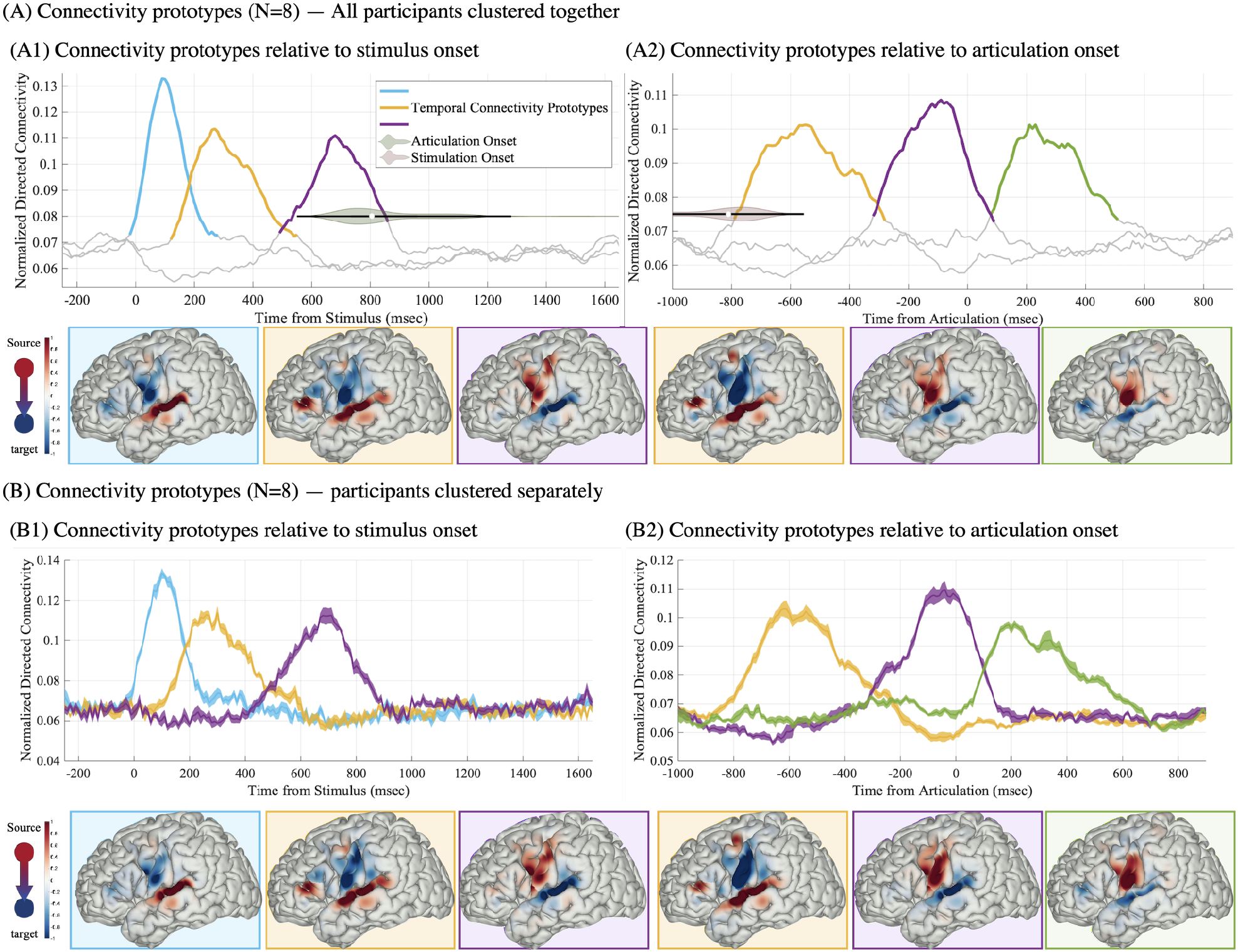
Connectivity prototypes in auditory repetition; variability across participants. (A) The connectivity prototypes and their corresponding source and target distributions when directed connectivity profiles from *N* =8 participants are clustered together (repeated from Fig. 3E and F); (B) The mean of the connectivity prototypes and corresponding average source and target distribution when directed connectivity profiles from *N* =8 participants are clustered separately. Shaded area around the curves in B shows standard error of the mean across participants.

### S.1 Passive listening as a negative control

Three of the patients in our cohort, performed a passive listening version of the auditory repetition task. In a separate recording block, participants were instructed to passively listen to identical stimuli from the auditory repetition task. We first provide the average neural activity across STG, IFG, pre-, and post-central cortices locked to stimulus presentation for both the passive listening (Fig. S5 A) and the auditory repetition (Fig. S5 B) tasks. In the passive listening task after stimulus presentation is over (range: 430 − 790 msec, mean: 583 msec from stimulus onset), there is a delay and the task moves to the next trial, while in the auditory repetition task articulation is engaged. For this reason, we focused our analysis on the interval [-250, 800] msec from stimulus onset (Fig. S5 C,D). This allows us to cover both stimulus presentation and pre-articulatory periods while avoiding the start of the next trials in the passive listening condition.

We first replicated the connectivity prototypes for the auditory repetition condition limited to the [-250, 800] time-interval, an analysis that is different in length and number of participants from the results in the main text (i.e. compare Fig. 3 E to Fig. S5 F). Specifically, we found that clustering with the same number of components (K=4) reveals two comprehension prototypes (blue and yellow) followed by a pre-articulatory prototype (purple). We then applied the same framework to the passive listening task, however we used three cluster components (K=3) as our analysis revealed that this was sufficient to represent the data (Fig. S5 E). The two prototypes associated with comprehension (Fig. S5 E, blue and yellow) were replicated in the passive condition in overall timing and spatial distribution. While we did not find a prearticulatory prototype during passive listening, we were concerned that it may be obscured or overlapping with the baseline/noise component (Fig. S5 E gray dashed line). To address this, we repeated clustering in the passive listening condition with four clusters (K=4). The clustering still revealed the first two comprehension prototypes (see yellow and blue boxes in Fig. S5 G) but the third cluster was diminished showing a distinctly silent topography compared with the pre-articulatory prototype in the auditory repetition condition, as the reviewer suggested (see purple and red in Fig. S5 G).

**Figure S5:**
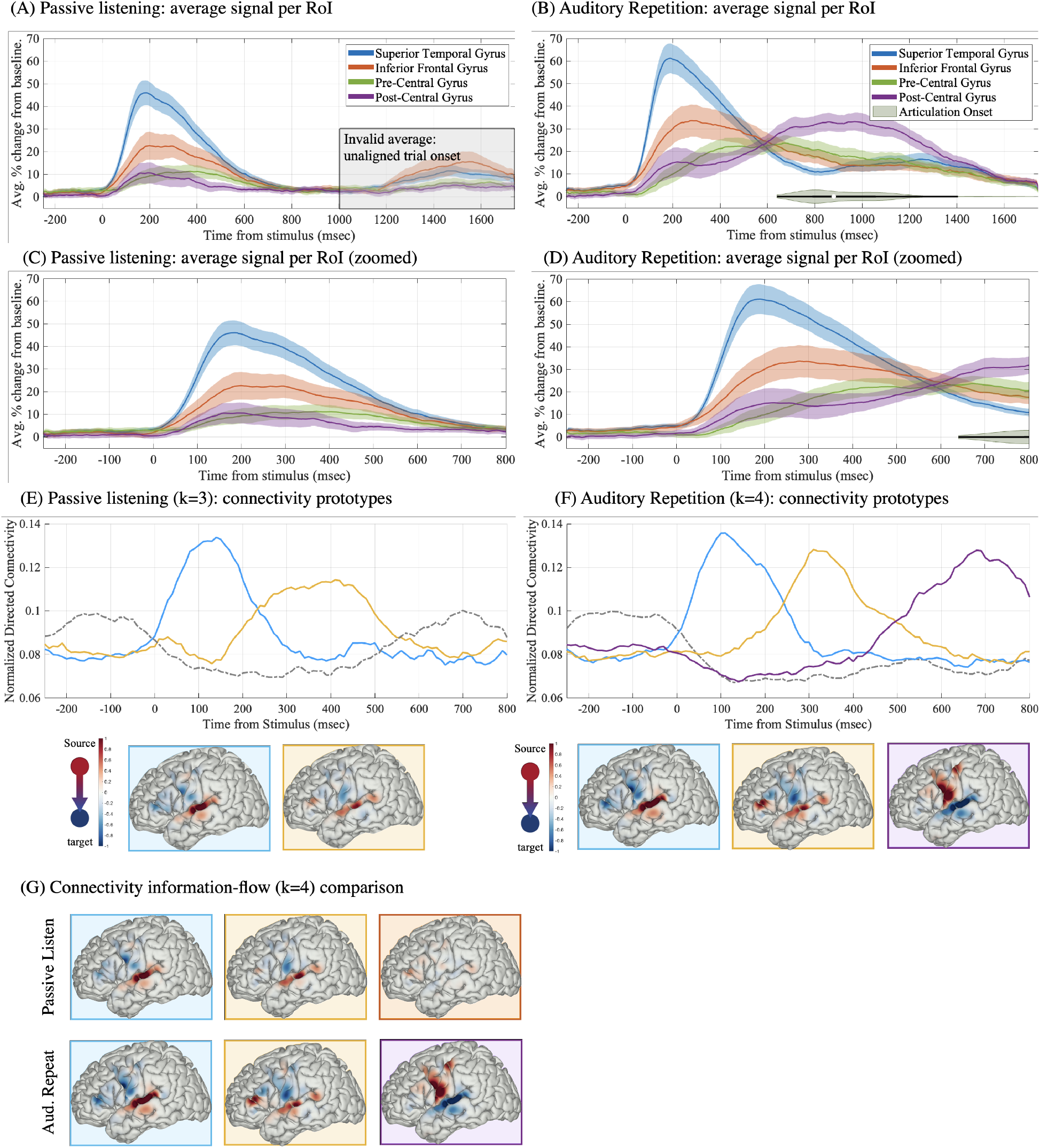
Passive listening as a control condition. Averaged neural activity in superior temporal, inferior frontal, pre-central, and post-central gyri are shown when trials are aligned to stimulus onset for (A) passively listening to auditory stimuli and (B) auditory repetition of the same stimuli. We focused our analysis on a time interval [-250, 800] that is comparable between the two conditions, which can be seen in C, D (a zoomed version of A and B, respectively). Results of directed connectivity modeling in the passive listening condition (E), and the repetition condition (F) are shown locked to stimulus (temporal prototypes and corresponding information sources and targets with *k*=3 and 4 cluster components, respectively). Regions of cortex showing information source (red) and target (blue) are colored for each prototype (the color of the box of each brain matches the color of the associated temporal curve in E and F). A comparison of the information sources and targets between the two conditions is shown in (G) applying the same number of of clusters in each analysis (*k*=4).

**Figure S6:**
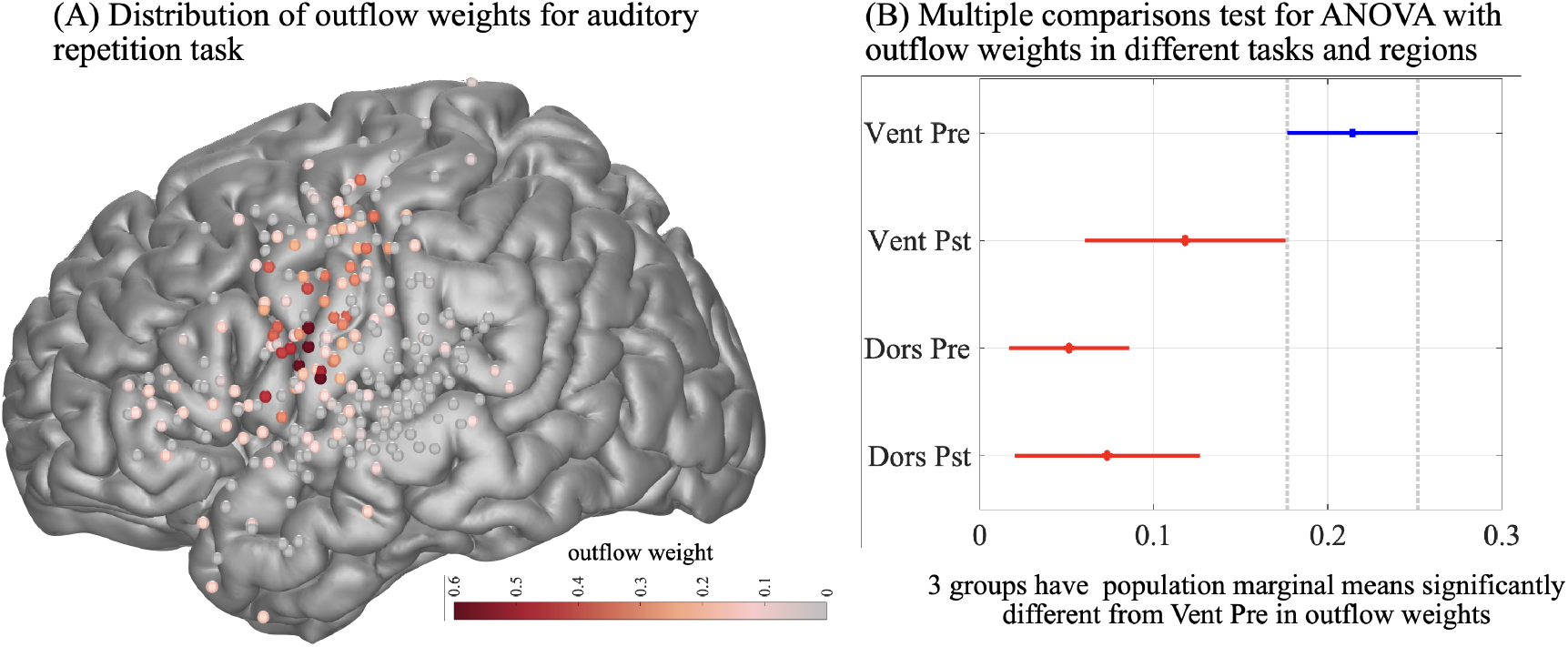
The corollary discharge source. (A) Spatial distribution of outflow weights associated with the corollary discharge prototype for the auditory repetition task. (B) Post-hoc multiple comparisons test using the result of ANOVA test for outflow weights with tasks and anatomical region (vental/dorsal pre- and post-central areas) as effects shows ventral pre-central gyrus as the main source of outflow associated with the corollary discharge prototype.

**Figure S7:**
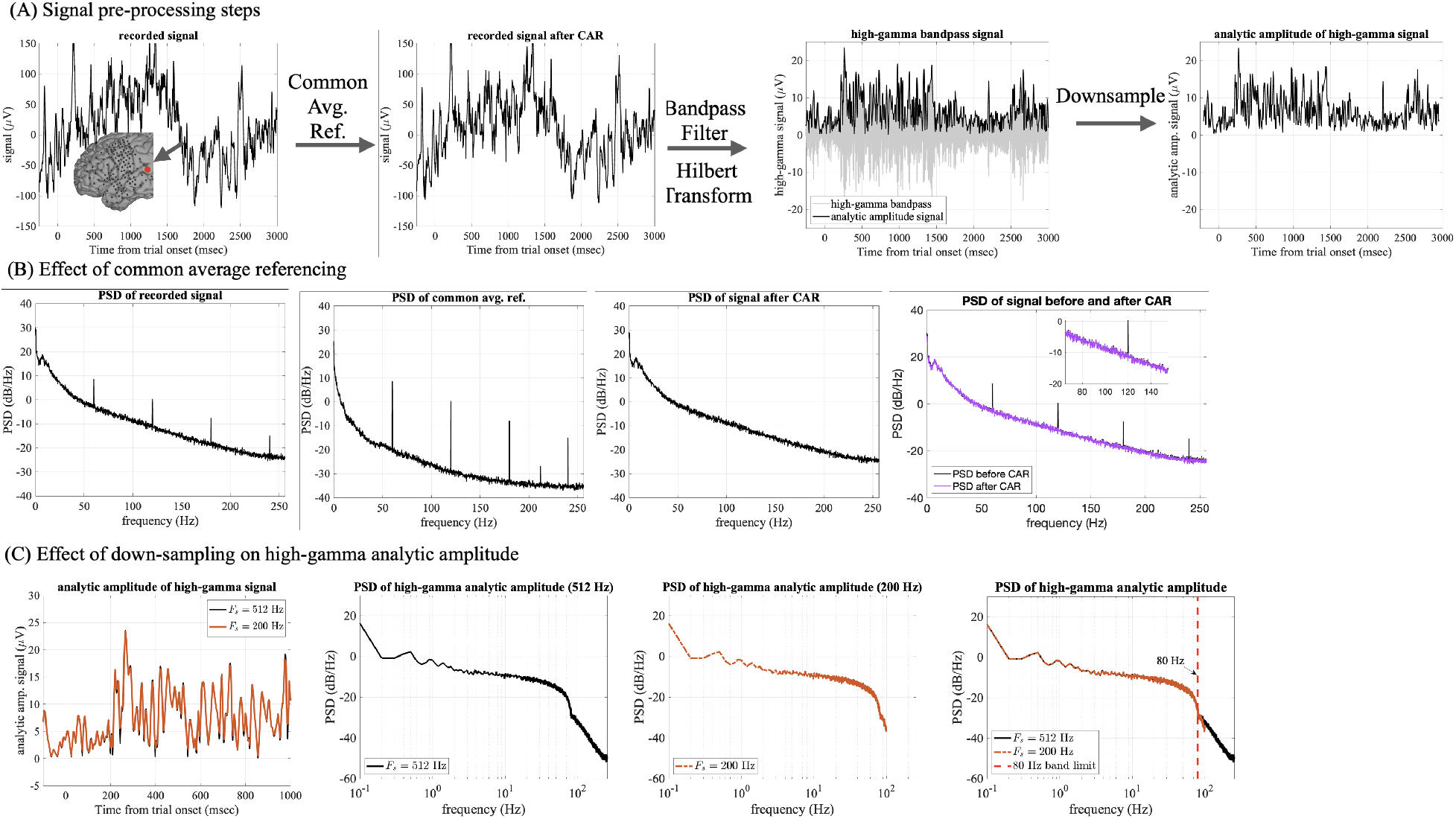
Illustration of the signal pre-processing stages. (A) The recorded signal is first referenced by subtracting the average of all electrodes at each time-point (common average reference), then band-pass filtered to 70-150 Hz high-gamma range and the analytic amplitude (envelope) of the high-gamma signal is extracted by a Hilbert transform. The resulting envelope signal is down-sampled from the original 512 Hz to 200 Hz. (B) The effect of the common average referencing is shown on the power spectral density of the signal. The major frequencies present in the common average signal are the 60 Hz line-noise and its harmonics, and by subtracting the effect of the line-noise is attenuated from the recorded signal. (C) The effect of down-sampling the analytic amplitude (envelope) of the high-gamma signal from 512 Hz to 200 Hz is shown in temporal domain (left plot) and frequency domain (power spectral density plots). High-gamma analytic amplitude is band-limited to 80 Hz and down-sampling to 200 Hz does not have an effect on the frequency content of the signal.

### S.2 Active electrode selection algorithm

Here, we introduce an unsupervised automatic algorithm to select a subset of electrodes that can be considered active given the ECoG data for a task. We expect active electrodes to have high event related responses, whereas the inactive ones to have a trial mean signals close to zero. We denote by *x*_*n*_(*r, t*) ∈ ℝ the ECoG signal for electrode *n* ∈ {1, *…, N* } at trial *r* ∈ {1, *…, R*} and time-step *t* ∈ {1, *…, T* }. We aim to find a subset of electrodes which have activity related to the task.

Motivated by this rationale, we first determined the trial mean signal for each electrode, i.e. 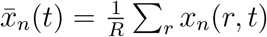. We empirically observed that further denoising this signal via wavelet-thresholding is beneficial. Let *W* denote the forward wavelet transform, *W*^*T*^ denote its inverse, and 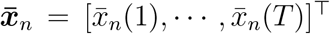. Let ℋ_*τ*_ (*x*) = *x* for |*x*| ≥ *τ* and 0 otherwise, be the hard-thresholding operator and similarly extend for vectors by applying element-wise. Then, the denoised mean signal can be represented by 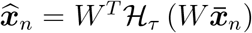 (see examples of the signals *x*_*n*_(*r, t*),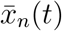, and 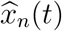 for three different electrodes in Fig. S8(a)). We used 5 levels of Daubechies 8 *(db8)* wavelet filters and we set *τ* = 0.5.

We computed the standard deviation of 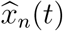 over time, i.e. 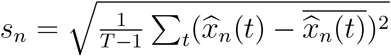.Higher values of *s*_*n*_ indicate active electrodes and smaller values indicate the inactive ones. We found a threshold *τ*_*s*_ such that the electrode *n* is considered active only if *s*_*n*_ *> τ*_*s*_ by fitting a Rayleigh-Rice mixture model (see a sample histogram of *s*_*n*_ and fitted model and threshold in Fig. S8(b)). The Rice component represented the active electrodes, while the Rayleigh component represented the inactive ones.

To describe this mixture model, let *α* indicate the probability that a sample *s* is from a Rayleigh distribution with parameter *b*^2^, and 1 − *α* the probability that *s* is from the Rice distribution with parameters *ν* and *σ*^2^. Then, the distribution of the mixture model can be written as

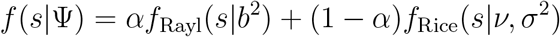

where Ψ = {*α, b*^2^, *ν, σ*^2^} and

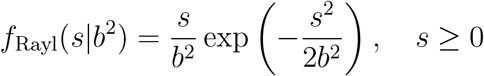

and

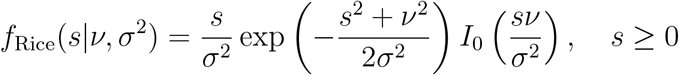

with *I*_0_(*·*) is the 0-th order modified Bessel function of first kind. Given the samples 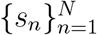 we found the parameters of this model (Ψ = {*α, b*^2^, *ν, σ*^2^}) via Expectation-Maximization (EM) algorithm for each participant (*61*). Given the fitted model parameters we set the threshold *τ*_*s*_ such that,

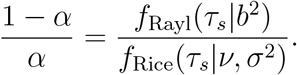

**Figure S8:**
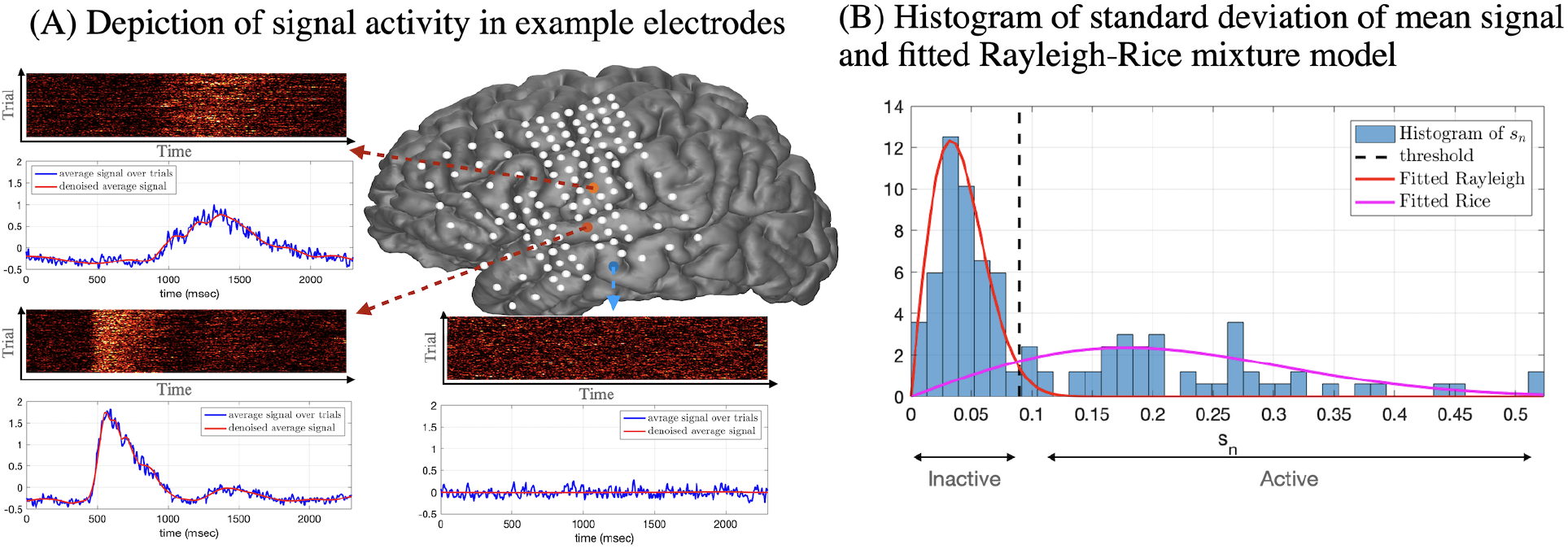
Example for active electrode selection algorithm. (A) Pictorial depiction of the signal *x*_*n*_(*r, t*), average signal over trials 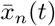, and denoised average signal 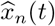 for two active and one inactive electrode. (B) Histogram of temporal changes from the mean, *s*_*n*_, for all the electrodes of one patient during auditory repetition task. Fitted Rayleigh-Rice mixture model and threshold are shown.

**Figure S9:**
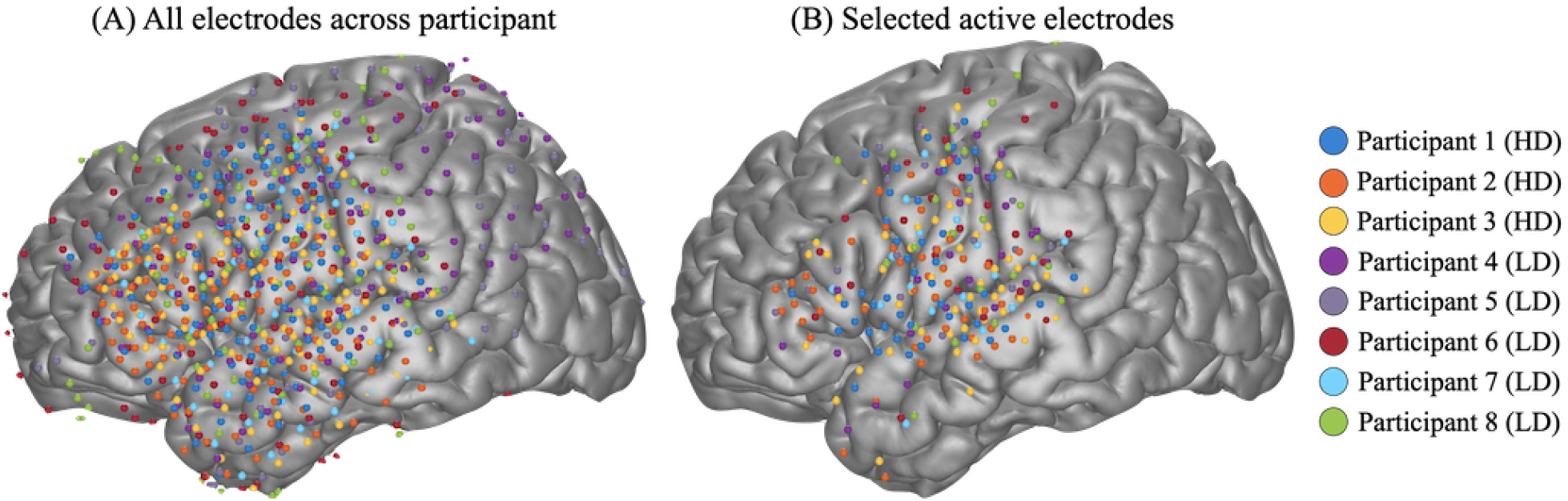
Electrode coverage and selection. (A) Implanted electrodes for all eight participants shown on a normal brain (MNI space, color-code represents participants). Five participants were implanted with low-density (LD) grids with 10mm spacing while three participants consented to be implanted with a hybrid-density (HD) grid with 10 mm overall spacing and 5mm spacing in specific areas. (B) Resulting electrodes from the active electrode selection algorithm are shown.

**Figure S10:**
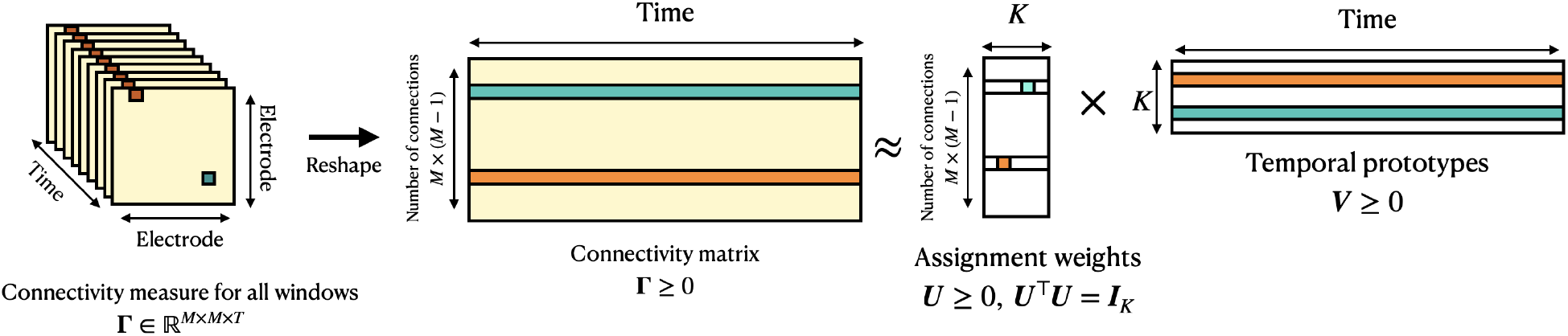
Unsupervised clustering applied to temporal connectivity profiles. A pictorial representation of the orthogonal non-negative matrix factorization (ONMF) algorithm applied to connectivity measures across time windows. The connectivity tensor (of size electrode x electrode x time) is reshaped into a matrix (of size connection-number x time). Temporal connectivity profiles (rows of the connectivity matrix **Γ**) are clustered into *K* prototypes (rows of the matrix **V**) with their corresponding assignment weights to a cluster (non-zero element in each row of the matrix **U**). Similar to other dimensionality reduction algorithms like principal component analysis (PCA), the connectivity matrix **Γ** is approximated by the lower-dimensional matrices **U** and **V**. Matrix factorization via ONMF, in contrast to PCA, assigns each connection to only one prototype (only one element in each row of **U** can be non-zero) and thus yields a clustering algorithm.

**Figure S11:**
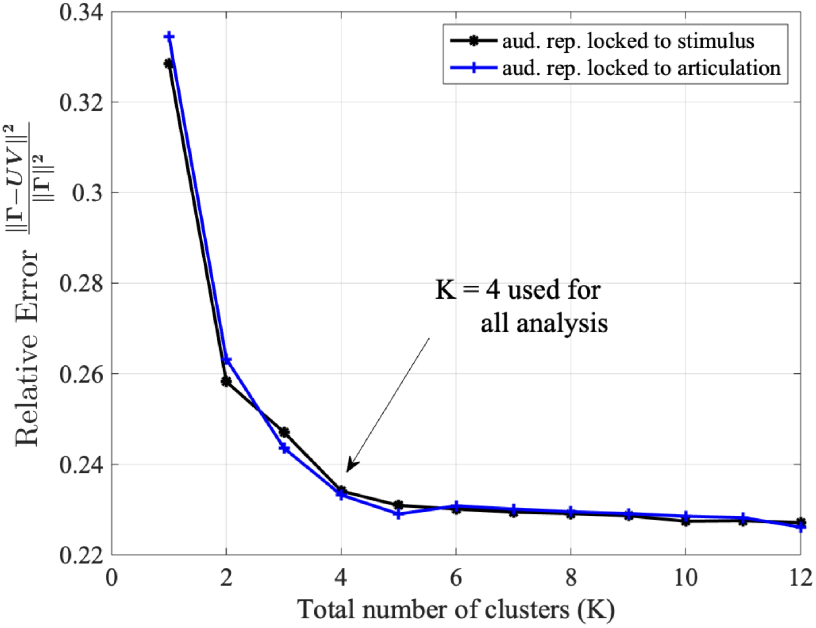
Number of clusters. Relative recovery error, 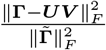, computed for different number of cluster components, *K*, and shown for auditory repetition task locked to stimulus (black) and articulation (blue). We choose *K* = 4 components in our analysis.

**Table S1:** Start time of consecutive activity (above 1.96 SD of baseline; longer than 100 msec) relative to stimulus onset across electrodes in different regions of interest during the auditory repetition task (mean *±* standard deviation across electrodes).

| RoI | STG | IFG | Pre-central | Post-central |
| --- | --- | --- | --- | --- |
| Time from stimulus (msec) | 105 ± 37 | 181 ± 42 | 435 ± 114 | 845 ± 53 |

